# Developmental maturation of millimeter-scale functional networks across brain areas

**DOI:** 10.1101/2024.05.28.595371

**Authors:** Nathaniel J. Powell, Bettina Hein, Deyue Kong, Jonas Elpelt, Haleigh N. Mulholland, Matthias Kaschube, Gordon B. Smith

**Author notes:** Corresponding author: Gordon Smith. Contributed equally (Nathaniel J. Powell, Bettina Hein). Co-supervised work (Matthias Kaschube, Gordon B. Smith). The authors declare no competing financial interests.

## Abstract

Interacting with the environment to process sensory information, generate perceptions, and shape behavior engages neural networks in brain areas with highly varied representations, ranging from unimodal sensory cortices to higher-order association areas. Recent work suggests a much greater degree of commonality across areas, with distributed and modular networks present in both sensory and non-sensory areas during early development. However, it is currently unknown whether this initially common modular structure undergoes an equally common developmental trajectory, or whether such a modular functional organization persists in some areas—such as primary visual cortex—but not others. Here we examine the development of network organization across diverse cortical regions in ferrets of both sexes using *in vivo* widefield calcium imaging of spontaneous activity. We find that all regions examined, including both primary sensory cortices (visual, auditory, and somatosensory—V1, A1, and S1, respectively) and higher order association areas (prefrontal and posterior parietal cortices) exhibit a largely similar pattern of changes over an approximately 3 week developmental period spanning eye opening and the transition to predominantly externally-driven sensory activity. We find that both a modular functional organization and millimeter-scale correlated networks remain present across all cortical areas examined. These networks weakened over development in most cortical areas, but strengthened in V1. Overall, the conserved maintenance of modular organization across different cortical areas suggests a common pathway of network refinement, and suggests that a modular organization—known to encode functional representations in visual areas—may be similarly engaged in highly diverse brain areas.

**Significance:** Different areas of the mature brain encode vastly different representations of the world. This study shows that a modular functional organization where nearby neurons participate in similar functional networks is shared across different brain areas not only during early development, but also as the brain matures where it remains a shared feature that shapes neural activity. The largely conserved trajectory of developmental changes across brain areas suggests that similar circuit mechanisms may drive this maturation. This implies that the large literature on developing cortical circuits, which is largely focused on sensory areas, may also apply more broadly, and that perturbations during development that impinge on any such shared mechanisms may produce deficits that extend across multiple brain systems.

## Introduction

In the mature cortex, neurons selective for features of the internal or external environment participate in highly interconnected neural networks that provide the basis for perception and behavior. Across a range of cortical areas, the information processed by these networks varies greatly, from primarily unimodal sensory representations in areas such as A1, V1, and S1, to more complex and higher order representations in areas such as PPC and PFC. The developmental origin of this functional diversity is a central question in cortical development, as is the relative balance of common organizational features versus early area-specific functions. Current evidence suggests that the specialization of cortical areas is thought to begin early in development, with genetic cues defining broad area identity, which is later refined through area-specific inputs in an activity dependent manner (Sur and Rubenstein, 2005; Cadwell et al., 2019), although the degree to which this cortical maturation occurs in concert across diverse brain areas remains unclear.

Much of what is known about the developmental maturation of functional properties at the network level comes from studies of primary sensory cortices. Here, the properties of the mature network emerge over the course of development through considerable refinement at both the neuron and network level. For example, in V1, from the time neurons can be driven by visual stimuli they become increasingly selective to stimulus features such as edge orientation (Chapman and Stryker, 1993) and direction of motion (Li et al., 2006). Here, these representations are organized into a modular structure, with nearby neurons exhibiting highly similar selectivity for a range of visual properties (Hubel and Wiesel, 1968; Blasdel and Salama, 1986; Weliky et al., 1996; Issa et al., 2000; Kara and Boyd, 2009; Smith et al., 2015b). Notably, once this modular representation of orientation preference is established around eye-opening, it remains stable across further development (Chapman et al., 1996). However, well before the onset of these sensory-evoked responses, functional neural networks exist in the visual cortex that are readily apparent through correlations in spontaneous activity (Chiu and Weliky, 2001). Activity in these early networks shows a highly dense and pronounced modular spatial structure, with nearby neurons showing highly coherent activity patterns that are organized into distributed functional networks comprised of multiple modules correlated across millimeters (Smith et al., 2018). Notably, these early correlated networks appear to be a precursor for the modular representation of orientation (Smith et al., 2018; Trägenap et al., 2023).

We recently found that during early development, such distributed and modular functional networks are not unique to V1, but are a common feature that is shared across diverse cortical areas. Both the primary sensory areas A1, V1, and S1, as well as the higher-order association areas PFC and PPC exhibit spontaneous activity that is strongly correlated amongst local populations, forming functional modules which themselves form patchy distributed networks (Powell et al., 2024). To date, these modular functional networks outside V1 have only been examined during early development, well before the onset of mature sensory-evoked responses and the key developmental transition to externally-driven sensory-evoked activity. Thus it remains unclear if the continued presence of modular functional organization seen in V1 is indicative of a visual-specific process, or rather is it a reflection of a more global and universal developmental program shared across the cortex.

To address this question, we used ongoing spontaneous activity to assess functional network organization across the cortex at two key periods in development: first, around the time of eye-opening and ear canal opening in the ferret (P27-32), which marks a major developmental milestone and a transition in the nature of incoming sensory information; and secondly 1-2 weeks later (P39-43), when response properties in V1 have been shown to approach mature levels (Li et al., 2006; Smith et al., 2015a). Examining spontaneous activity allows us to measure and compare intrinsic network organization across areas, without the need to design optimal stimuli for both sensory and non-sensory areas. Comparing this data to that which we previously obtained from P21-24 (Powell et al., 2024), we find that the developmental maturation of network properties is highly conserved across cortical areas. Across all developmental ages examined, we find that spontaneous activity remained significantly modular in both sensory and non-sensory cortical areas. All areas exhibited a similar weakening of modular organization, while also continuing to show distributed and modular correlations across millimeters. Correlation strength decreased and activity became higher dimensional in all areas except V1, which showed increasing millimeter-scale correlations with age. Together, these results demonstrate that the modular functional organization that serves as a common developmental origin for highly diverse cortical areas (Powell et al., 2024) remains a common feature across different cortical areas which undergo a largely common developmental trajectory.

## Materials and Methods

### Data collection

#### Animals

All experimental procedures were approved by the University of Minnesota Institutional Animal Care and Use Committee and were performed in accordance with guidelines from the US National Institutes of Health. We obtained 27 male and female ferret kits from Marshall Farms and housed them with jills on a 16-h light/8-h dark cycle. No statistical methods were used to predetermine sample sizes, but our sample sizes are similar to those reported in previous publications.

#### Viral Injection

Viral injections were performed as previously described (Smith & Fitzpatrick, 2016). Briefly, we expressed GCaMP6s (Chen et al., 2013) in neurons by microinjecting AAV1.hSyn.GCaMP6s.WPRE.SV40 (Addgene) into layer 2/3 of targeted cortical areas at P13-32, 7-21 d before imaging. Anesthesia was induced with isoflurane (4–5%) and maintained with isoflurane (1– 1.5%). Glycopyrolate (0.01 mg/kg) or Atropine (0.02 mg/kg) and bupivacaine/lidocaine (1:1 mixture) were administered, and animal temperature was maintained at approximately 37 °C with a water pump heat therapy pad (Adroit Medical HTP-1500, Parkland Scientific). Animals were mechanically ventilated and both heart rate and end-tidal CO_2_ were monitored throughout the surgery (Digicare LifeWindow). Using aseptic surgical technique, skin and muscle overlying target areas were retracted, and a small burr hole was made with a handheld drill (Fordom Electric Co.). Approximately 1 μL of virus contained in a pulled-glass pipette was pressure injected into the cortex at two or three depths (∼200 - 400 μm below the surface) over 20 min using a Nanoject-III (World Precision Instruments). The craniotomy was sealed and the skin sutured closed.

Targets for different cortical areas were as follows:

V1: ∼6-8 mm lateral from midline, ∼1-2 mm anterior to the Lamda

PPC: ∼1-2 mm lateral from midline, ∼4 mm posterior to Bregma

A1: ∼7-9 mm lateral to midline, ∼3 mm posterior to Bregma

S1: ∼2-3 mm lateral to midline, ∼ 1 mm anterior to Bregma

PFC: ∼1-2 mm lateral to midline, ∼7-8 mm anterior to Bregma

We injected virus into and imaged 1-3 areas per animal. In most experiments, multiple areas were labeled and imaged in a single animal (mean 1.9 ± 0.2 areas per animal).

#### Cranial window surgery

On the day of experimental imaging, ferrets age P27-43 were anesthetized with 3%–4% isoflurane and atropine (0.2 mg/kg) or glycopyrrolate (0.01 mg/kg) was administered. Animals were placed on a feedback-controlled heating pad to maintain an internal temperature of 37 °C. Animals were intubated and ventilated. Isoflurane was delivered between 1 and 2% throughout the surgical procedure to maintain a surgical plane of anesthesia. An intraperitoneal or intravenous catheter was placed to deliver fluids. EKG, end tidal CO_2_, and internal temperature were continuously monitored during the procedure and subsequent imaging session. The scalp was retracted and a custom titanium headmount adhered to the skull using C&B Metabond (Parkell). A 6 to 7 mm craniotomy was performed over areas of viral expression and the dura was retracted to reveal the cortex. Cover glass (round, #1.5 thickness, Electron Microscopy Sciences) adhered to the bottom of a custom titanium or 3-D printed plastic insert was placed onto the brain to gently compress the underlying cortex and dampen biological motion during imaging. Upon completion of the surgical procedure, isoflurane was gradually reduced (0.6 to 1.0%) and then vecuronium bromide (2 mg/kg/hr) was delivered to reduce motion and prevent spontaneous respiration.

#### Widefield epifluorescence and two-photon imaging

Spontaneous activity was recorded in a quiet darkened room for 10-40 minutes per area. For consistency with prior results (Smith et al., 2018; Powell et al., 2024) we performed our experiments under light isoflurane anesthesia, which has been shown in V1 to not disrupt the spatial structure of modular functional activity (Smith et al., 2018). Widefield epifluorescence imaging was performed with a Zyla 5.5 sCMOS camera (Andor) controlled by MicroManager software (Edelstein et al., 2014). Images were acquired at 15 Hz with 4 × 4 binning to yield 640 × 540 pixels.

#### Histology

Following imaging, animals were euthanized with 5% Isoflurane and pentobarbital. Animals were perfused with heparinized saline solution followed by 4% paraformaldehyde, then the brains were removed and kept for histology. Viral expression of GCaMP was documented in the intact brain with appropriate excitation and emission filters. Images of expression were aligned to a common coordinate system using prominent brain features (sulci, fissures, and external edges of brain). Areas of expression from each brain were outlined manually in Matlab, and are shown in Fig. 1.

**Figure 1:**
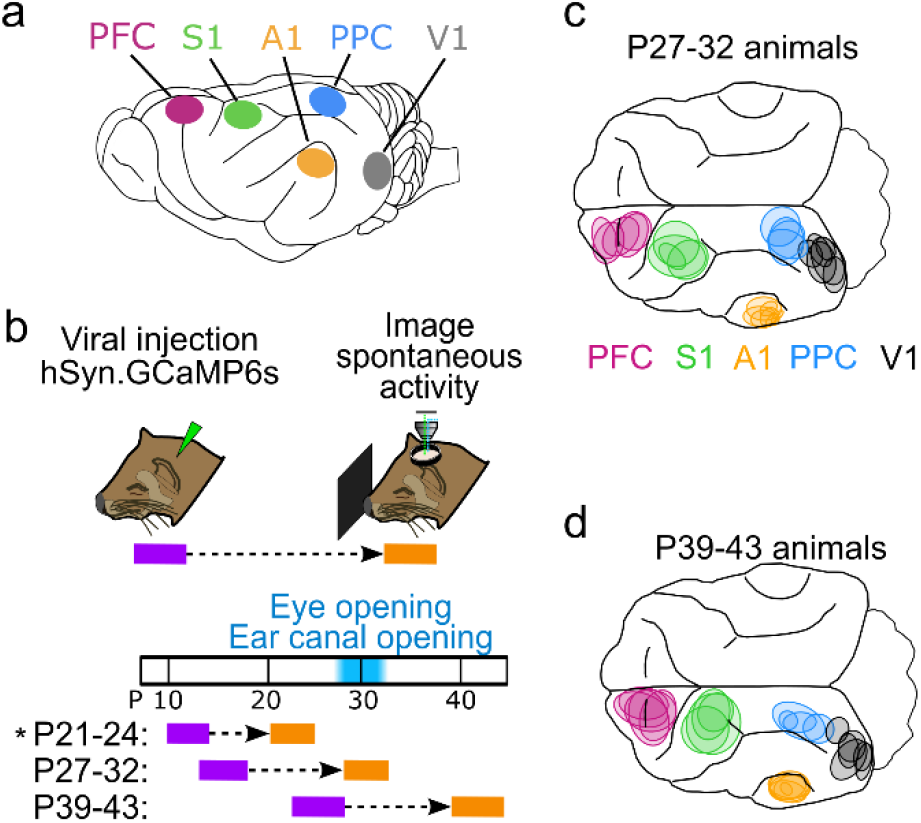
Imaging distinct cortical regions in ferrets at different developmental ages. **a**. Target cortical locations. PFC – prefrontal cortex, S1 – somatosensory cortex, A1 – auditory cortex, PPC – posterior parietal cortex, V1 – visual cortex. **b**. Experimental timeline. Animals were injected with AAV expressing GCaMP6s 10-14 days prior to imaging. Imaging was performed at P27-32 or P39-43. P21-24 data from ref (Powell et al., 2024) (indicated by *) is presented for comparison with other ages. **c**. Imaged locations for P27-32 animals reconstructed from histology, colored based on assigned cortical area. **d**. Same as (c) for P39-43 animals.

### Analysis methods

#### Widefield data pre-processing

Widefield data pre-processing, event extraction and calculation of spontaneous correlations was performed largely as described (Smith et al., 2018) for all imaged areas. Briefly, to correct for mild brain movement during imaging, we registered each imaging frame by maximizing phase correlation to a common reference frame. A region of interest (ROI) was manually drawn around the cortical area with high and robust spontaneous activity. ROIs were also drawn to remove any artifacts or debris in the visible field of view (FOV). The baseline fluorescence (F_0_) for each pixel was obtained by applying a median filter to the raw fluorescence trace with a window between 10 and 64 seconds. Filter width was chosen for each imaging session individually, such that the baseline followed faithfully the slow trend of the fluorescence activity. The baseline corrected activity was calculated as

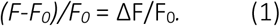

#### Event detection

To detect spontaneously active events, we first determined active pixels on each frame using a pixel-wise threshold set to 3 standard deviations above each pixel’s mean value across time. Active pixels not part of a contiguous active region of at least 0.01mm^2^ were considered ‘inactive’ for the purpose of event detection in order to minimize detecting noise as ‘active’ pixels. Active frames were taken as frames with a spatially extended pattern of activity (>40% of pixels were active).

Consecutive active frames were combined into a single event starting with the first high activity frame and then either ending with the last high activity frame or, if present, an activity frame defining a local minimum in the fluorescence activity. In order to assess the spatial pattern of an event, we extracted the maximally active frame for each event (the “event frame”), which was defined as the frame with the highest activity averaged across the ROI.

#### Calculation of correlation patterns

To assess the spatial correlation structure of spontaneous or evoked activity, we applied a Gaussian spatial band-pass filter (with SD of Gaussian filter kernel *s*_high_=195μm, s_low_=26-41μm) to each event frame and down-sampled it to 160 x 135 pixels. The resulting patterns, named *A*_*i*_ in the following, where i=1,…,N, were used to compute the spontaneous correlation patterns as the pairwise Pearson’s correlation between all locations ***x*** within the ROI and the seed point ***s***

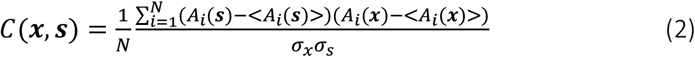

Here the brackets < > denote the average over all *N* patterns and *σ*_***x***_ denotes the standard deviation of *A* over all *N* patterns at location ***x***.

#### Shuffled control ensemble and surrogate correlation patterns

To evaluate the statistical significance of quantities characterizing the correlation patterns observed during spontaneous activity, we compared the real ensemble of spontaneous activity patterns from a given experiment with a control ensemble, obtained by eliminating most of the spatial relationships between the patterns. To this end, all activity patterns were randomly rotated (rotation angle drawn from a uniform distribution between 0° and 360° with a step size of 10°) and reflected (with probability 0.5, independently at the x- and y-axis at the center of the ROI), resulting in an equally large control ensemble with similar statistical properties, but little systematic interrelation between patterns. Surrogate correlation patterns were then computed from these ensembles as described above.

#### Dimensionality of spontaneous activity

We estimated the cross-validated dimensionality *d*_eff_ of the subspace spanned by spontaneous activity patterns (see (Stringer et al., 2019)). First, we randomly divided the activity patterns into two non-overlapping subsets X_1_ and X_2_, and then performed PCA on X_1_ to find the axis spanned by it. Next, we projected X_2_ onto these PCs to estimate the variance λ_i_ explained by each PC. Lastly, dimensionality was calculated as (Abbott et al., 2011):

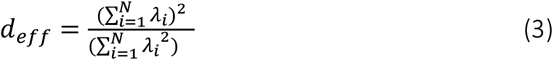

#### Spatial range of correlations

To assess the strength of spontaneous correlations over distance and compare across experiments with different numbers of spontaneous events, we computed the variance of pixelwise correlation values located at a given distance from a seed point (Dahmen et al., 2022; Powell et al., 2024). Closer to the seed point, correlations will have both strongly positive and strongly negative values, leading to a high variance. In contrast, further away from the seed point correlations will be closer to zero and exhibit reduced variance. To assess long-range correlation strength, we computed the variance within a ring from 1.8 – 2.2 mm from seed point. To control for the finite number of events in each experiment, we also computed the variance for surrogate correlation patterns (see above) generated for each experiment using a matched number of patterns. Subtracting this control variance for each experiment allows for comparison across experiments with varying numbers of observed events. The ROI for two A1 FOVs were too small to compute a surrogate dataset at 2mm, and were excluded from this analysis.

#### Modularity and wavelength estimation

To estimate the wavelength of individual calcium events, we calculated the 1-D radial average of the spatial autocorrelation of the band-pass-filtered activity pattern. The wavelength of the event was taken as twice the distance to the first minimum from the origin. Modularity is a measure of the regularity of the spatial arrangement of activity domains within the pattern. The modularity of each event was calculated as the absolute difference in amplitude between the first minimum and the subsequent maximum of the 1-D radial averaged autocorrelation.

To determine if the modularity observed during spontaneous events was statistically significant, we compared it to a distribution of modularity values for inactive frames used as the control. Control frames were drawn from the bottom 10% of frames lacking an identified spontaneous event (see above) based on mean activity within the ROI. For each experiment, we obtained 100 sets of control frames containing an event-matched number of frames and calculated the median modularity across these frames, generating a distribution of 100 control modularity values. This was then compared to the median modularity across spontaneous events to obtain a p-value.

#### Module amplitude

We defined the module amplitude of an individual widefield event as the average amplitude (in ΔF/F, prior to spatial filtering) of the module peaks divided by the background activity. Peaks in activity for each event were first obtained by using the FastPeakFind.m function in Matlab (Adi, 2021). Background activity was taken as the median amplitude (in ΔF/F) of activity in locations ½ wavelength from the peak. Here, the average wavelength across all events was used for each FOV. The module amplitude of an event was taken as the average amplitude across all peaks in the event.

### Statistical analysis

Nonparametric statistical analyses were used throughout the study. All tests were two-sided unless otherwise noted. Mack-Skillings tests were used to test for global effects of age across all areas. Comparisons across age within areas were performed using a Kruskal-Wallis test.

Data analysis was performed in Matlab (Mathworks), and Python.

### Data, Materials, and Software Availability

All data and code used in this study are available upon request from the corresponding author.

## Results

### Distributed and modular spontaneous activity patterns in diverse cortical areas 10-14 days beyond eye-opening

We first sought to examine animals aged P27-32, which in the ferret is around the key developmental milestones of eye- and ear canal-opening (Figure 1, a-c). To assess the state of functional networks at this point in development we expressed the calcium sensor GCaMP6s and examined ongoing spontaneous activity through widefield calcium imaging in five different cortical areas. In both primary sensory (V1, A1, and S1) and association areas (PFC and PPC), we observed frequent spontaneous events that exhibited clear coordination in activity amongst local populations of neurons (Figure 2a,b). In all cortical areas, these spontaneous events exhibited clearly modular activity, with distinct patches of activity that were distributed across the cortical surface. Notably this modular structure was evident in unprocessed images without spatial filtering (Figure 2b,c *left*).

**Figure 2:**
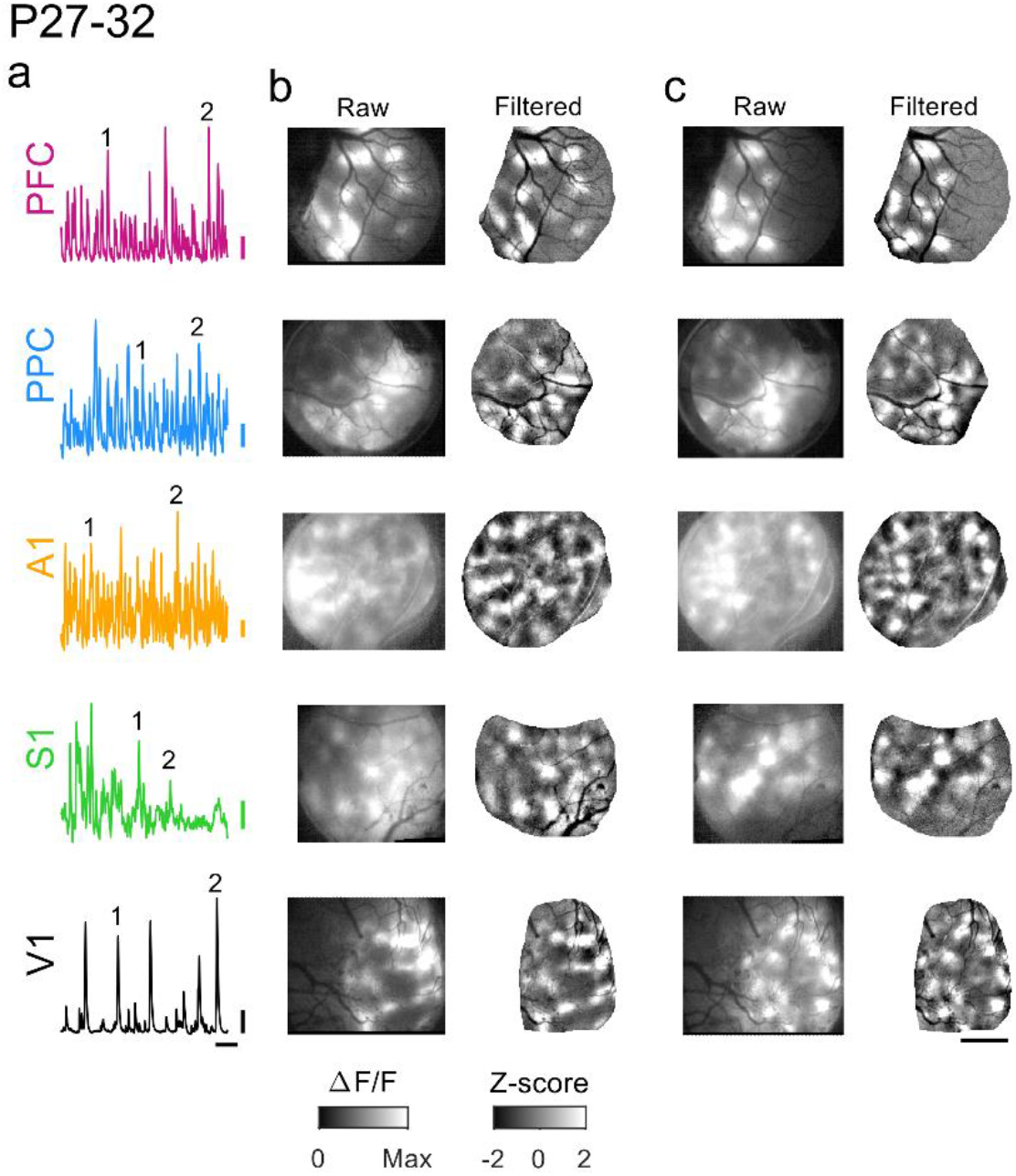
Spontaneous events exhibit widespread and distributed modular activity in diverse brain areas at P27-32. **a**. Example timecourse of spontaneous activity in different cortical areas. Numbers indicate events shown in b and c. **b**. Spontaneous events show modular patterns of activation in PFC, PPC, A1, S1, and V1 at P27-32. *Left*: raw event pattern showing clear modular patterns of activity in all areas at time (1) in (a). *Right*: Same event after applying a highpass spatial filter. **c**. Second representative event from same experiments as (b), at time (2) in (a). Scale bars (a): 0.1 ΔF/F, 20 sec; (b-c): 1 mm.

We next sought to determine whether similarly locally coordinated activity also existed across different cortical areas in animals one to two weeks past eye opening and ear canal opening (P39-43, Figure 1b,d). Strikingly, in these animals we observed that activity in spontaneous events continued to exhibit clear and pronounced modular structure, with patchy activity distributed across millimeters of cortex (Figure 3). As in younger animals, the modular structure in spontaneous events was evident in both spatially filtered images (Figure 3, b-c, *right*), as well as unprocessed images (Figure 3, b-c, *left*). The overall structure of spontaneous activity patterns was highly similar across areas and ages, and was also similar in appearance to the activity patterns seen previously in animals aged P21-24, 7-10 days before eye opening (Powell et al., 2024).

**Figure 3:**
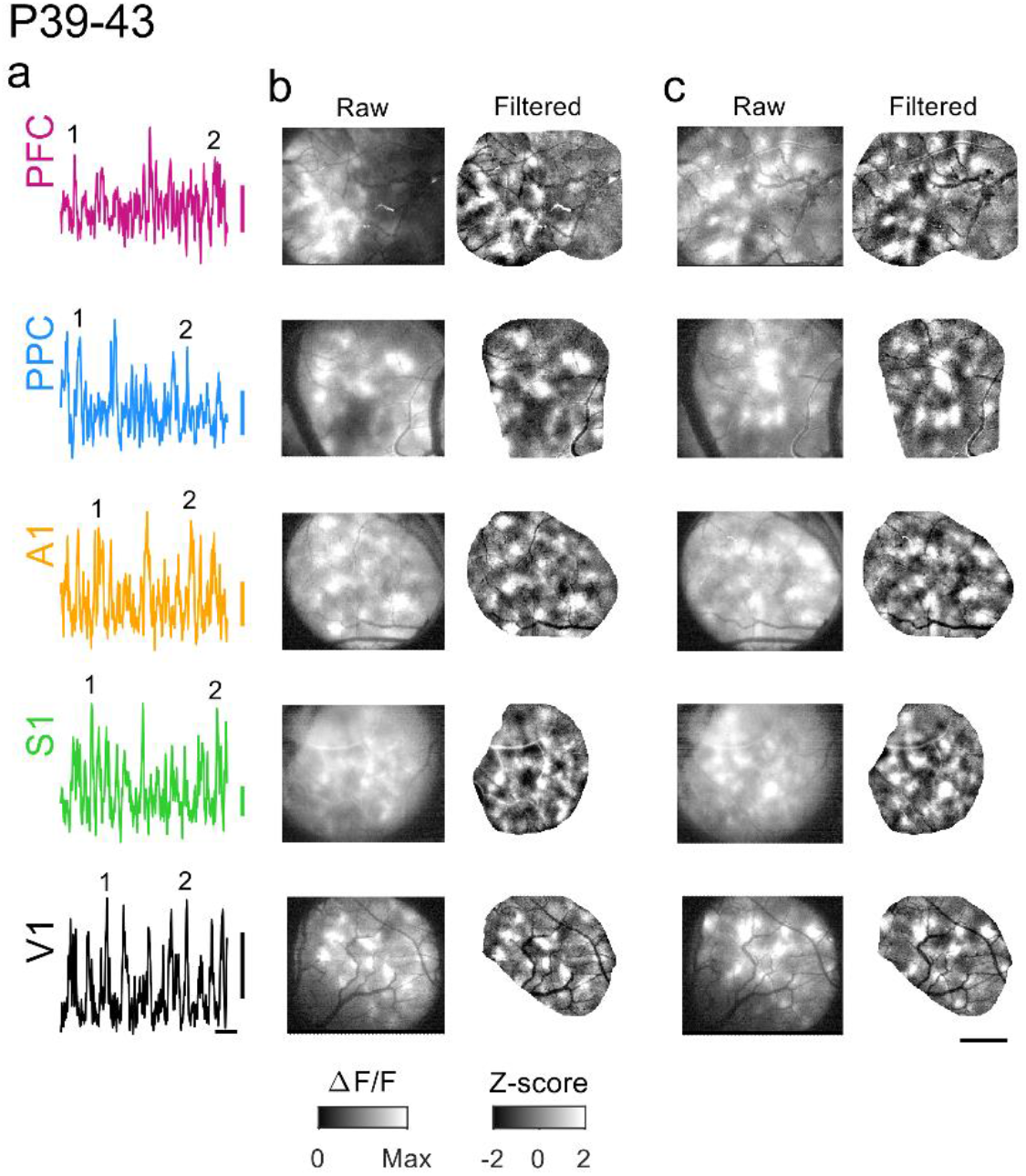
Spontaneous events continue to exhibit widespread and distributed modular activity at P39-43. **a**. Example timecourse of spontaneous activity in different cortical areas. Numbers indicate events shown in b and c. **b**. Spontaneous events show modular patterns of activation in all areas at P39-43. *Left*: raw event pattern showing clear modular patterns of activity in all areas at time (1) in (a). *Right*: Same event after applying a highpass spatial filter. **c**. Second representative event from same experiments as (b), at time (2) in (a). Scale bars (a): 0.1 ΔF/F, 20 sec; (b-c): 1 mm.

### Quantitative similarity in developmental trajectory of modular spontaneous activity

We next sought to directly compare the strength of modular activity in different brain areas over development. To achieve this, we calculated the modularity of each spontaneous event by first computing the spatial autocorrelation and then measuring the relative amplitude of the first trough and subsequent peak, providing a measure of the regularity in spacing of active domains (Powell et al., 2024). Comparing this data to spontaneous activity at P21-24 (Powell et al., 2024), the strength of this modularity showed a global decline with age across areas (Figure 4a, Mack-Skillings test for age while controlling for area, TS(2) = 26.73, p < 0.001), with significant declines within area for PFC, PPC and S1 (Kruskal Wallis p<0.05 for each, see Table 4-1). Notably however, events in all cortical areas at both P27-32 and P39-43 remained significantly modular when compared to controls (26 of 26 vs shuffle at P27-32, 26 of 26 significant at P39-43 at p<0.05 in all cases).

**Figure 4:**
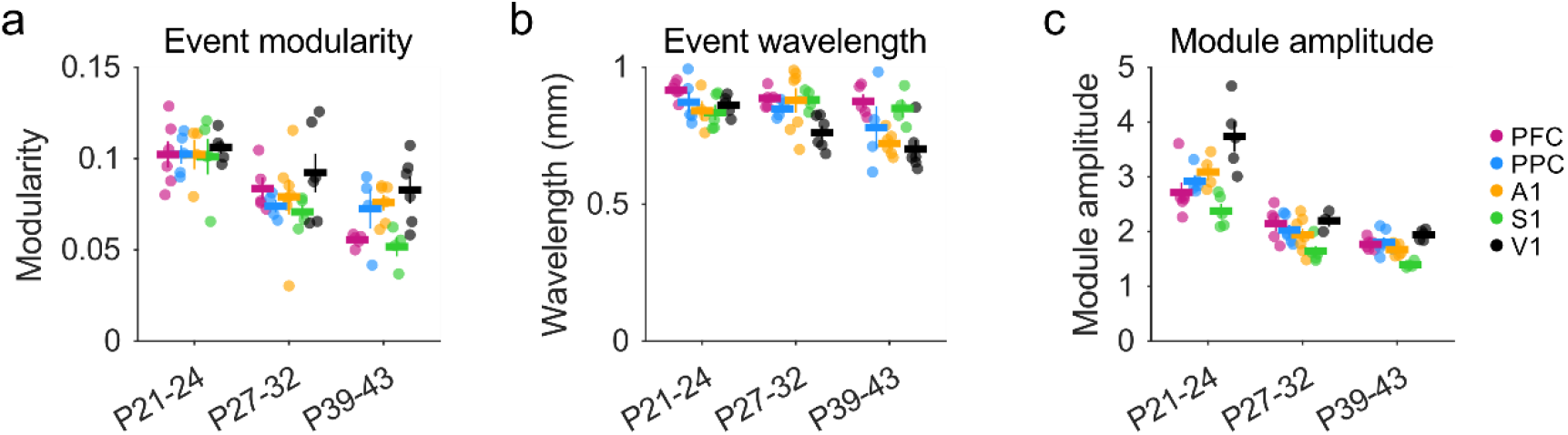
Developmental maturation of modular spontaneous activity across cortical areas. **a**. Modularity of spontaneous events declines with age, but remains significant in all cases versus shuffle controls. **b**. The wavelength of active modules is relatively consistent across areas and shows a slight but significant decline with age in A1. **c**. The amplitude of active modules declines in a similar manner with age in all cortical areas examined. P21-24 data in (a-c) replotted for comparison from (Powell et al., 2024).

We next measured the spatial wavelength of active domains, again using the spatial autocorrelation of spontaneous events. Overall, the wavelength of activity decreased slightly with age (Figure 4b), which was significant across areas (Mack-Skillings test for age while controlling for area, TS(2)= 12.31 p = 0.002), however within each area, these changes were only significant in A1 (Kruskal Wallis, p<0.05, see Table 4-2). Notably, the wavelength of spontaneous activity in all cortical areas at all ages is roughly similar to that of orientation preference maps in V1

The modularity of events derived from spatial autocorrelations primarily reflects the degree of regularity in the spacing of active domains and is less sensitive to how strongly modules are active relative to surrounding cortex. We therefore computed the ‘module amplitude’ to assess the selectivity of modular activation (see Methods), and observed a corresponding decline with age across areas (Figure 4c, Mack-Skillings test for age while controlling for area, TS(2) = 46.51, p< 0.001; with PPC, PFC, S1 and V1 showing significant within-area declines, Kruskal Wallis p<0.05 for each, see Statistics Table 4-3). Together, these results indicate that cortical networks across multiple brain regions continue to exhibit large-scale patterns of distributed modular spontaneous activity well beyond the onset of sensory-evoked responses, and that these diverse cortical areas undergo a similar trajectory of developmental maturation.

### Modular activity shows millimeter-scale correlations in all areas through P39-43

During early development, spontaneous activity not only shows modular patterns of activity, but also millimeter-scale correlations (Smith et al., 2018; Powell et al., 2024). The patterns of spontaneous activity at these ages comprise a low dimensional subset of all theoretically possible activity patterns, such that certain sets of modules tend to be co-active across events, giving rise to millimeter scale correlations in activity in all cortical areas examined (Powell et al., 2024). We next sought to determine if the observed weakening of modular activity (Figure 4a,c) is accompanied by a corresponding diversification in millimeter-scale networks, which could result from activity that is locally modular but undergoes a developmental loss in larger-scale organization. Alternatively, large-scale correlated networks could remain present in diverse cortical areas, consistent with multiple studies have found that millimeter-scale modular networks remain a prominent feature in V1 in the mature brain (Kenet et al., 2003; O’Hashi et al., 2018; Smith et al., 2018). In order to determine if the large-scale network structure that is common in the early cortex (Powell et al., 2024) follows distinct developmental trajectories in V1 versus other cortical areas, we computed correlations across all spontaneous events observed in widefield imaging.

We found that in all cortical areas examined, at P27-32, spontaneous activity continued to show long-range correlations, with strongly correlated modules frequently appearing separated by millimeters of cortical area (Figure 5a). Notably, multiple functional networks were reflected in the structure of these correlations, as evidenced by the diversity in the spatial pattern of correlations for different seed points (Figure 5a, *left* vs. *right*). When we examined correlations at P39-43, we observed that the overall structure remained roughly similar in all cortical areas, with positively correlated modules separated by several millimeters. As in younger ages, different cortical locations participated in differently correlated networks (Figure 5b, *left* vs. *right*). However, the strength of these correlations appeared to decrease with age in all areas other than V1, with correlation patterns showing generally weaker and less distinct structure than at earlier ages. This progression was qualitatively similar across areas outside of V1, suggesting a similar developmental trajectory across these regions.

**Figure 5:**
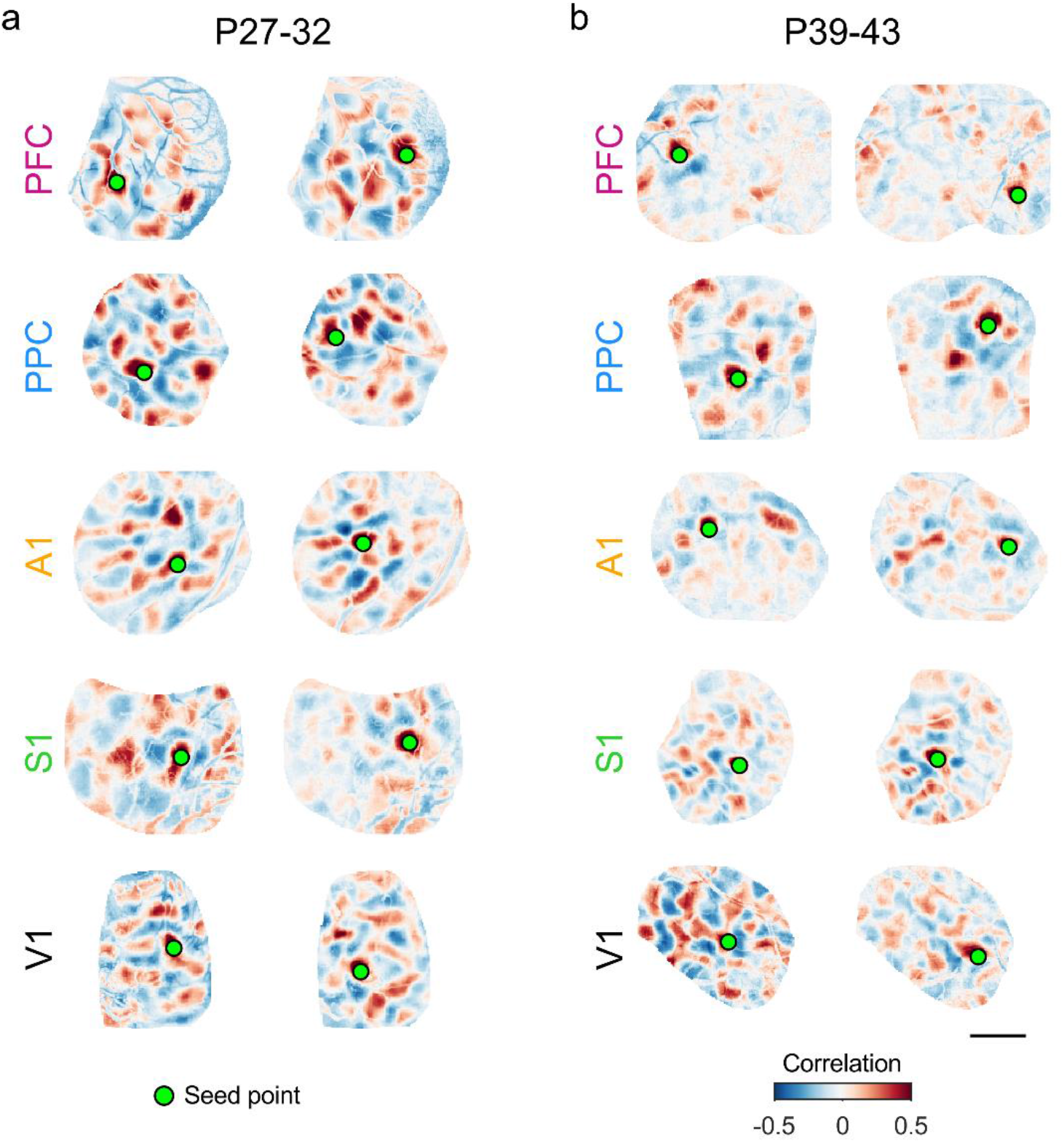
Millimeter-scale modular correlations across diverse brain areas after eye-opening. **a**. Correlations across spontaneous events for representative experiments at P27-32. Pixelwise correlations are shown relative to two different seed points for each area. Note that the spatial patterns of correlations vary between seed points, reflecting multiple distinct correlated networks. **b**. Modular correlations are still present at P39-43 in all cortical areas examined. Scale bar: 1 mm.

### Developmental maturation in millimeter-scale correlated networks across brain areas

We next sought to quantify the strength of these long-range correlations and their developmental progression across areas. Given that the absolute amplitude of correlations can be influenced by the finite number of spontaneous events we are able to observe in each experiment, making direct comparisons of correlation amplitude across areas and ages is challenging. To address this, we employed a measure of correlation strength that allows for the correction of spurious correlations which result from finite event numbers. Thus, correlation strength was determined computing the variance in pixel-wise correlations (Dahmen et al., 2022; Powell et al., 2024) at a distance of 2 mm (bin size 1.8-2.2 mm) from the seed point. Strong correlations will exhibit high variance (as they include both strongly positive and strongly negative peaks), whereas weak correlations will show low variance. Importantly, this approach allows for subtracting the contribution of artifacts in correlation strength resulting from finite event numbers, which we assessed using surrogate correlation patterns (see methods). In nearly all cases at both P27-32 and P39-43, the strength of these long-range correlations was statistically significant relative to shuffled controls (23 of 25 vs shuffle at P27-32, 23 of 25 significant at P39-43 at p<0.05). A critical advantage of this approach is that it allows comparison across experiments at different ages and areas with different numbers of observed events. When comparing our data to that from animals at P21-24 published previously (Powell et al., 2024), we find that correlation variance at 2 mm changes significantly with age across areas (Figure 6a, Mack-Skillings test for age while controlling for area, TS(2) = 22.91, P < 0.001). Within areas, PFC, PPC, S1 and A1, exhibited significant declines with age (Kruskal Wallis test, p< 0.05 for all areas, see Statistics Table 6-1). In S1, the largest decline occurred between P21-24 and P27-32, whereas PFC, PPC and S1 only showed large declines by P39-43. In contrast to the other brain regions examined, correlation variance in V1 showed significant increases across age (Kruskal Wallis test, p=0.047, see Statistics Table 6-1). Together these results show that despite developmental changes in strength that exhibit some variation across brain areas, cortical networks continue to exhibit significant long-range correlations at P39-43.

**Figure 6:**
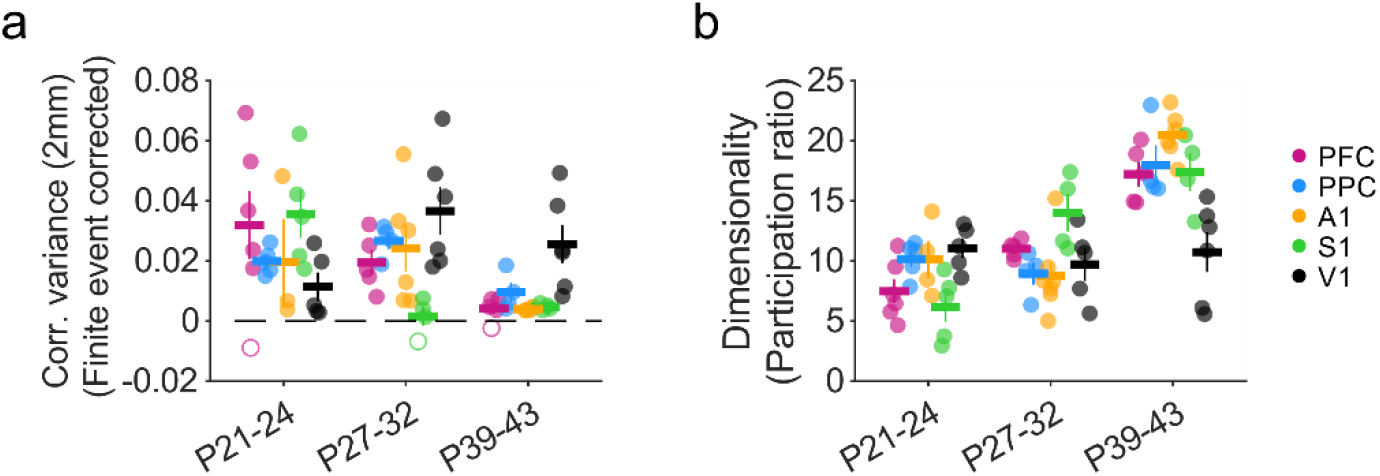
Maturation of correlation strength and dimensionality across cortical areas. **a**. Long-range correlations (1.8-2.2 mm) remain statistically significant versus control across age in all areas, while showing a significant decline with age in all areas except V1. Closed circles indicate experiments with significant correlations relative to shuffled controls. Open circles are non-significant at p<0.05. **b**. Dimensionality increases significantly with age in most cortical areas. Data from P21-24 animals in (a-c) replotted for comparison from (Powell et al., 202

Spontaneous activity at early developmental timepoints (P21-24) is characterized by activity patterns that reside in a relatively small subspace of all possible activity patterns, which can be described with a relatively low number of dimensions (Powell et al., 2024). In order to determine whether the complexity and diversity of spontaneous activity changed with age, we therefore computed the cross-validated dimensionality (Abbott et al., 2011; Stringer et al., 2019) of widefield spontaneous activity for each experiment. We observed significant changes in the dimensionality of activity across areas (Figure 6b, Mack-Skillings test for age while controlling for area, TS(2) = 33.08, P < 0.001). Dimensionality increased significantly in PFC, PPC, and A1 (Kruskal Wallis test, p< 0.05 for all areas, see Statistics Table 6-2), while remaining moderately low compared to the space of all possible activity patterns. S1 showed a similar increase in dimensionality to these areas, but was not statistically significant (Kruskal Wallis test, p=0.14, see Statistics Table 6-2). In contrast, the dimensionality of activity in V1 appeared to change very little with age (Kruskal Wallis test, p=0.77, see Statistics Table 6-2). Consistent with the decrease in correlation strength that we observe with age, we saw an increase in dimensionality with development in PFC, PPC, S1, and A1, that was absent in V1. The continued low-dimensionality and strengthening long-range correlations in V1 may be an area-specific specialization, potentially reflecting the contribution of the highly organized long-range horizontal projections that develop following eye-opening (Gilbert and Wiesel, 1989; Callaway and Katz, 1990; Durack and Katz, 1996; Bosking et al., 1997). Collectively, these results show that millimeter-scale correlations and moderately low dimensional activity remain a central feature of spontaneous activity over development in both sensory and non-sensory cortices.

## Discussion

By examining multiple cortical areas serving diverse functions—ranging from primary sensory cortices to higher-order association areas—at different points in development, we found evidence for a common modular structure in spontaneous activity that undergoes a largely common developmental maturation. Our data build upon prior results which show that across multiple brain areas in early development before to the transition to sensory-evoked activity, local circuits exhibit highly dense and modular activity that is coherent on both the local and millimeter-scale (Powell et al., 2024).The results presented here show that this common organization in the early cortex is followed by a largely conserved developmental trajectory across many diverse cortical areas including both sensory and non-sensory cortices. Initially low dimensional and strongly correlated activity becomes increasingly rich and high dimensional with age, while retaining clear locally modular organization and long-range correlations over a period spanning major developmental milestones of eye-opening and ear canal opening. Notably, our results show that the central features of these networks—modular organization and millimeter-scale correlations—are conserved across cortical areas and ages, persisting well after this developmental transition to predominantly extrinsically driven activity. Over development, quantitative differences in correlation strength and dimensionality emerge, primarily between V1 and other areas, which reflect the interplay between a common functional architecture and area-specific specializations.

Our finding of a modular functional organization across many diverse cortical areas outside of early development is consistent with several prior observations. In mature V1, the presence of a modular functional organization is well documented in the structure of visually-evoked feature maps (Hubel and Wiesel, 1968; Blasdel and Salama, 1986; Weliky et al., 1996; Issa et al., 2000; Kara and Boyd, 2009; Smith et al., 2015b). This modular organization is also present in spontaneous activity (Kenet et al., 2003; O’Hashi et al., 2018; Smith et al., 2018), which we also see in our results. Likewise, our finding that such distributed modular correlated networks are present in other sensory areas (A1, S1) well after the transition to extrinsically-driven activity is consistent with the presence of modular organization of feature representations reported in these areas (Sur et al., 1981; Recanzone et al., 1999; Schreiner et al., 2000; Friedman et al., 2004; Atencio and Schreiner, 2012). Furthermore, in the mature PFC, anatomical projections corresponding to distinct information streams show spatial clustering (Goldman-Rakic and Schwartz, 1982; Kritzer and Goldman-Rakic, 1995). Whether the correlated networks we observe in spontaneous activity in these areas correspond to such representations—as they do in V1(Kenet et al., 2003; O’Hashi et al., 2018; Smith et al., 2018)—remains to be determined.

While our results in general reflect a fairly consistent developmental maturation across highly varied cortical areas, there are several notable differences between brain regions in our data. Amongst several different metrics, activity in S1 shows large changes between P21-24 and P27-32, which appear to occur later (between P27-32 and P39-43) in other brain areas. These results suggest that the timing of maturation may differ across cortical areas, with S1 progressing faster than other areas. This is consistent with the relatively earlier onset of sensory evoked and thalamocortically-driven responses in S1 relative to V1 (Huberman et al., 2008; Antón-Bolaños et al., 2019). The diversity of brain regions is generally thought to initially arise from broad genetically-specified gradients that are subsequently refined in an activity-dependent manner driven by area-specific feed-forward inputs (Sur and Rubenstein, 2005; Cadwell et al., 2019). In this context, the relatively stronger modularity and millimeter-scale correlations seen in V1 compared to other cortical areas in older animals may reflect the emergence of a strong alignment of low-dimensional feed-forward input—encoding selectivity for edge orientation—with locally modular intracortical networks (Trägenap et al., 2023).

Prior work has suggested that in early development, when intracortical horizontal connections are thought to be primarily short-range (Durack and Katz, 1996), recurrent local circuits can give rise to distributed and modular correlations through self-organizing mechanisms (Smith et al., 2018; Mulholland et al., 2024). In such theoretical models (von der Malsburg, 1973; Ernst et al., 2001), this modular structure arises through the interaction of local excitation and lateral inhibition (LE/LI), and predicts a tight coupling of activity in excitatory and inhibitory populations, which has been observed in V1 in early development (Mulholland et al., 2021). Such LE/LI mechanisms appear to be engaged in early V1 where unstructured optogenetic activation gives rise to modular cortical activity (Mulholland et al., 2024). It is possible that a similar mechanism may operate in early development in other cortical regions, although this remains to be tested. In addition, direct optogenetic stimulation of the cortex across early to late developmental periods could be used to tease apart the relative contributions of inputs to these different brain regions, and determine if the changes we observe with age are due a developmental shift towards input-driven activity and the cortex becoming less strongly dominated by recurrent circuits.

In V1, long-range horizontal axons emerge over development and exhibit distributed and patchy arborizations that align spatially with functional modules representing stimulus features (Gilbert and Wiesel, 1989; Callaway and Katz, 1990; Bosking et al., 1997). Similar patchy horizontal axonal projections have been seen in a wide range of cortical areas, and have been suggested to reflect a canonical wiring motif in the mature cortex (Douglas and Martin, 2004; Muir and Douglas, 2011). If such structured long-range connections emerge over development as a common feature across brain regions, they could serve to constrain and stabilize the large-scale correlated networks we observe across areas in spontaneous activity in multiple cortical areas, thereby maintaining this millimeter-scale organization in the face of local sparsification and increased dimensionality. It is also possible that differences in the strength of such specific long-range horizontal connections across areas might also contribute to the relatively stronger millimeter-scale correlations seen in V1 at P39-43.

The question of whether functional modules such as those we observe in correlated spontaneous activity play a role in cortical computations has long been the subject of investigation. Modular organization can lead to efficient networks by minimizing connection distances between functionally coupled neurons (Koulakov and Chklovskii, 2001), and can also lead to computational advantages (Meunier et al., 2010). Our finding that modular networks in most brain regions weaken and become higher dimensional with age suggests that such an organization may play a developmental role in coarsely grouping neurons into functional units in early development before giving way to more locally-diverse organization. Whether the modules we observe in different cortical areas in animals well past eye-opening reflect a persistent feature of developmental network organization, or rather contribute directly to cortical computation, remains to be determined and could potentially result in different answers across different cortical areas.

## Acknowledgments

We would like to thank Casey Xamonthiene, Nic Glewwe, Matt Paruzynski, Jack Kapler, and Sophie Bowman for experimental support. We would like to thank Sigrid Trägenap and Lorenzo Butti for analysis support. We would also like to thank members of the Smith and Kaschube labs for helpful discussions. We thank Len White for inspiring discussions encouraging this project.

This work was supported by these funding sources:

National Eye Institute (NIH R01EY030893-01)

(GBS) Bundesministerium für Bildung und Forschung (BMBF 01GQ2002)

(MK) Whitehall Foundation (2018-05-57)

(GBS) National Science Foundation (IIS-2011542) (GBS)

## Author contributions

GBS and MK conceptualized the study. NJP and GBS performed the surgeries and collected the data. NJP, BH and JE performed the pre-processing on the data. BH, GBS, and MK designed the analyses and metrics. All authors analyzed the data and contributed to the figures. GBS and MK acquired funding and supervised the project. GBS wrote the original draft. All authors reviewed and edited the final text.

---

**Table 4-1:** Event modularity mean (sem)

| Area | P21 - 24 | P27 - 32 | P39 - 43 | Kruskall Wallis across age |
| --- | --- | --- | --- | --- |
| PFC | 0.102 (0.007), n=6 | 0.084 (0.006), n=5 | 0.055 (0.001), n=5 | **p = 0.003, H(2) = 11.32** |
| PPC | 0.102 (0.005), n=5 | 0.074 (0.004), n=4 | 0.072 (0.011), n=4 | **p = 0.014, H(2) = 8.54** |
| A1 | 0.102 (0.008), n=4 | 0.079 (0.010), n=7 | 0.076 (0.004), n=6 | p = 0.619, H(2) = 0.96 |
| S1 | 0.101 (0.010), n=5 | 0.071 (0.004), n=4 | 0.052 (0.005), n=4 | **p = 0.013, H(2) = 8.72** |
| V1 | 0.106 (0.004), n=5 | 0.092 (0.011), n=6 | 0.083 (0.008), n=6 | p = 0.459, H(2) = 1.56 |

**Table 4-2:** Event wavelength mean (sem)

| Area | P21 - 24 | P27 - 32 | P39 - 43 | Kruskall Wallis across age |
| --- | --- | --- | --- | --- |
| PFC | 0.917 (0.013), n=6 | 0.887 (0.015), n=5 | 0.876 (0.025), n=5 | p = 0.210, H(2) = 3.12 |
| PPC | 0.873 (0.037), n=5 | 0.848 (0.017), n=4 | 0.780 (0.078), n=4 | p = 0.368, H(2) = 2.00 |
| A1 | 0.841 (0.036), n=4 | 0.879 (0.045), n=7 | 0.722 (0.018), n=6 | **p = 0.034, H(2) = 6.79** |
| S1 | 0.835 (0.029), n=5 | 0.882 (0.019), n=4 | 0.850 (0.032), n=4 | p = 0.724, H(2) = 0.65 |
| V1 | 0.862 (0.016), n=5 | 0.762 (0.025), n=6 | 0.701 (0.033), n=6 | p = 0.096, H(2) = 4.68 |

**Table 4-3:** Module amplitude mean (sem)

| Area | P21 - 24 | P27 - 32 | P39 - 43 | Kruskall Wallis across age |
| --- | --- | --- | --- | --- |
| PFC | 2.72 (0.19), n=6 | 2.15 (0.14), n=5 | 1.77 (0.04), n=6 | **p = 0.003, H(2) = 11.81** |
| PPC | 2.92 (0.10), n=5 | 2.03 (0.10), n=6 | 1.81 (0.09), n=6 | **p = 0.050, H(2) = 6.00** |
| A1 | 3.09 (0.16), n=4 | 1.94 (0.12), n=7 | 1.68 (0.05), n=6 | p = 0.343, H(2) = 2.14 |
| S1 | 2.38 (0.13), n=5 | 1.64 (0.09), n=5 | 1.40 (0.03), n=4 | **p = 0.002, H(2) = 12.02** |
| V1 | 3.74 (0.28), n=5 | 2.20 (0.11), n=3 | 1.94 (0.04), n=5 | **p = 0.009, H(2) = 9.40** |

**Table 6-1:** Correlation strength - 2mm mean (sem)

| Area | P21 - 24 | P27 - 32 | P39 - 43 | Kruskall Wallis across age |
| --- | --- | --- | --- | --- |
| PFC | 0.031 (0.012), n=6 | 0.019 (0.005), n=5 | 0.003 (0.002), n=5 | **p = 0.004, H(2) = 11.02** |
| PPC | 0.014 (0.003), n=5 | 0.026 (0.004), n=4 | 0.015 (0.006), n=4 | **p = 0.042, H(2) = 6.32** |
| A1 | 0.022 (0.018), n=3 | 0.027 (0.007), n=6 | 0.004 (0.000), n=5 | **p = 0.014, H(2) = 8.55** |
| S1 | 0.033 (0.005), n=5 | 0.002 (0.003), n=4 | 0.005 (0.001), n=4 | **p = 0.006, H(2) = 10.11** |
| V1 | 0.011 (0.006), n=5 | 0.036 (0.007), n=6 | 0.023 (0.007), n=6 | **p = 0.047, H(2) = 6.12** |

**Table 6-2:** Dimensionality mean (sem)

| Area | P21 - 24 | P27 - 32 | P39 - 43 | Kruskall Wallis across age |
| --- | --- | --- | --- | --- |
| PFC | 7.48 (1.01), n=6 | 11.01 (0.32), n=5 | 17.23 (1.04), n=5 | **p = 0.002, H(2) = 12.04** |
| PPC | 10.17 (0.66), n=5 | 8.95 (0.92), n=4 | 17.96 (1.68), n=4 | **p = 0.005, H(2) = 10.50** |
| A1 | 10.13 (1.54), n=4 | 8.76 (1.19), n=7 | 20.47 (0.79), n=6 | **p = 0.020, H(2) = 7.86** |
| S1 | 6.16 (1.21), n=5 | 14.03 (1.58), n=4 | 17.38 (1.57), n=4 | p = 0.144, H(2) = 3.88 |
| V1 | 11.07 (0.83), n=5 | 9.72 (1.37), n=5 | 10.74 (1.66), n=6 | p = 0.777, H(2) = 0.50 |

## References

Abbott LF, Rajan K, Sompolinsky H (2011) Interactions between Intrinsic and Stimulus-Evoked Activity in Recurrent Neural Networks. In: The Dynamic Brain: An Exploration of Neuronal Variability and Its Functional Significance (Glanzman MD and D, ed), pp 65–82. New York: Oxford University Press, Inc.

Adi N (2021) Fast 2D peak finder. MATLAB Central File Exchange Available at: https://www.mathworks.com/matlabcentral/fileexchange/37388-fast-2d-peak-finder.

Antón-Bolaños N, Sempere-Ferràndez A, Guillamón-Vivancos T, Martini FJ, Pérez-Saiz L, Gezelius H, Filipchuk A, Valdeolmillos M, López-Bendito G (2019) Prenatal activity from thalamic neurons governs the emergence of functional cortical maps in mice. Science 364:987–990.

Atencio CA, Schreiner CE (2012) Spectrotemporal Processing in Spectral Tuning Modules of Cat Primary Auditory Cortex. PLOS ONE 7:e31537.

Blasdel GG, Salama G (1986) Voltage-sensitive dyes reveal a modular organization in monkey striate cortex. Nature 321:579–585.

Bosking WH, Zhang Y, Schofield B, Fitzpatrick D (1997) Orientation selectivity and the arrangement of horizontal connections in tree shrew striate cortex. J Neurosci 17:2112–2127.

Cadwell CR, Bhaduri A, Mostajo-Radji MA, Keefe MG, Nowakowski TJ (2019) Development and Arealization of the Cerebral Cortex. Neuron 103:980–1004.

Callaway EM, Katz LC (1990) Emergence and refinement of clustered horizontal connections in cat striate cortex. Journal of Neuroscience 10:1134–1153.

Chapman B, Stryker MP (1993) Development of orientation selectivity in ferret visual cortex and effects of deprivation. The Journal of neuroscience : the official journal of the Society for Neuroscience 13:5251–5262.

Chapman B, Stryker MP, Bonhoeffer T (1996) Development of orientation preference maps in ferret primary visual cortex. The Journal of neuroscience : the official journal of the Society for Neuroscience 16:6443–6453.

Chen TW, Wardill TJ, Sun Y, Pulver SR, Renninger SL, Baohan A, Schreiter ER, Kerr RA, Orger MB, Jayaraman V, Looger LL, Svoboda K, Kim DS (2013) Ultrasensitive fluorescent proteins for imaging neuronal activity. Nature 499:295–300.

Chiu C, Weliky M (2001) Spontaneous activity in developing ferret visual cortex in vivo. J Neurosci 21:8906–8914.

Dahmen D, Layer M, Deutz L, Dąbrowska PA, Voges N, von Papen M, Brochier T, Riehle A, Diesmann M, Grün S, Helias M (2022) Global organization of neuronal activity only requires unstructured local connectivity Palmer SE, Behrens TE, eds. eLife 11:e68422.

Douglas RJ, Martin KA (2004) Neuronal circuits of the neocortex. Annu Rev Neurosci 27:419–451.

Durack JC, Katz LC (1996) Development of horizontal projections in layer 2/3 of ferret visual cortex. Cereb Cortex 6:178–183.

Edelstein AD, Tsuchida MA, Amodaj N, Pinkard H, Vale RD, Stuurman N (2014) Advanced methods of microscope control using μManager software. J Biol Methods 1:e10.

Ernst UA, Pawelzik KR, Sahar-Pikielny C, Tsodyks MV (2001) Intracortical origin of visual maps. Nat Neurosci 4:431–436.

Friedman RM, Chen LM, Roe AW (2004) Modality maps within primate somatosensory cortex. Proceedings of the National Academy of Sciences 101:12724–12729.

Gilbert CD, Wiesel TN (1989) Columnar specificity of intrinsic horizontal and corticocortical connections in cat visual cortex. J Neurosci 9:2432–2442.

Goldman-Rakic PS, Schwartz ML (1982) Interdigitation of contralateral and ipsilateral columnar projections to frontal association cortex in primates. Science 216:755–757.

Hubel DH, Wiesel TN (1968) Receptive fields and functional architecture of monkey striate cortex. The Journal of physiology 195:215–243.

Huberman AD, Feller MB, Chapman B (2008) Mechanisms underlying development of visual maps and receptive fields. Annual review of neuroscience 31:479–509.

Issa NP, Trepel C, Stryker MP (2000) Spatial frequency maps in cat visual cortex. The Journal of neuroscience : the official journal of the Society for Neuroscience 20:8504–8514.

Kara P, Boyd JD (2009) A micro-architecture for binocular disparity and ocular dominance in visual cortex. Nature 458:627–631.

Kenet T, Bibitchkov D, Tsodyks M, Grinvald A, Arieli A (2003) Spontaneously emerging cortical representations of visual attributes. Nature 425:954–956.

Koulakov AA, Chklovskii DB (2001) Orientation Preference Patterns in Mammalian Visual Cortex: A Wire Length Minimization Approach. Neuron 29:519–527.

Kritzer MF, Goldman-Rakic PS (1995) Intrinsic circuit organization of the major layers and sublayers of the dorsolateral prefrontal cortex in the rhesus monkey. Journal of Comparative Neurology 359:131–143.

Li Y, Fitzpatrick D, White LE (2006) The development of direction selectivity in ferret visual cortex requires early visual experience. Nature neuroscience 9:676–681.

Meunier D, Lambiotte R, Bullmore E (2010) Modular and Hierarchically Modular Organization of Brain Networks. Frontiers in Neuroscience 4 Available at: https://www.frontiersin.org/articles/10.3389/fnins.2010.00200 [Accessed October 9, 2023].

Muir DR, Douglas RJ (2011) From Neural Arbors to Daisies. Cerebral Cortex 21:1118–1133.

Mulholland HN, Hein B, Kaschube M, Smith GB (2021) Tightly coupled inhibitory and excitatory functional networks in the developing primary visual cortex. Elife 10:e72456.

Mulholland HN, Kaschube M, Smith GB (2024) Self-organization of modular activity in immature cortical networks. :2024.03.02.583133 Available at: https://www.biorxiv.org/content/10.1101/2024.03.02.583133v1 [Accessed March 4, 2024].

O’Hashi K, Fekete T, Deneux T, Hildesheim R, van Leeuwen C, Grinvald A (2018) Interhemispheric Synchrony of Spontaneous Cortical States at the Cortical Column Level. Cereb Cortex 28:1794– 1807.

Powell NJ, Hein B, Kong D, Elpelt J, Mulholland HN, Kaschube M, Smith GB (2024) Common modular architecture across diverse cortical areas in early development. Proceedings of the National Academy of Sciences 121:e2313743121.

Recanzone GH, Schreiner CE, Sutter ML, Beitel RE, Merzenich MM (1999) Functional organization of spectral receptive fields in the primary auditory cortex of the owl monkey. Journal of Comparative Neurology 415:460–481.

Schreiner CE, Read HL, Sutter ML (2000) Modular Organization of Frequency Integration in Primary Auditory Cortex. Annual Review of Neuroscience 23:501–529.

Smith GB, Hein B, Whitney DE, Fitzpatrick D, Kaschube M (2018) Distributed network interactions and their emergence in developing neocortex. Nat Neurosci 21:1600–1608.

Smith GB, Sederberg A, Elyada YM, Van Hooser SD, Kaschube M, Fitzpatrick D (2015a) The development of cortical circuits for motion discrimination. Nature neuroscience 18:252–261.

Smith GB, Whitney DE, Fitzpatrick D (2015b) Modular Representation of Luminance Polarity in the Superficial Layers of Primary Visual Cortex. Neuron 88:805–818.

Stringer C, Pachitariu M, Steinmetz N, Carandini M, Harris KD (2019) High-dimensional geometry of population responses in visual cortex. Nature 571:361–365.

Sur M, Rubenstein JLR (2005) Patterning and Plasticity of the Cerebral Cortex. Science 310:805–810.

Sur M, Wall JT, Kaas JH (1981) Modular Segregation of Functional Cell Classes Within the Postcentral Somatosensory Cortex of Monkeys. Science 212:1059–1061.

Trägenap S, Whitney DE, Fitzpatrick D, Kaschube M (2023) The nature-nurture transform underlying the emergence of reliable cortical representations. :2022.11.14.516507 Available at: https://www.biorxiv.org/content/10.1101/2022.11.14.516507v2 [Accessed November 10, 2023].

von der Malsburg Chr (1973) Self-organization of orientation sensitive cells in the striate cortex. Kybernetik 14:85–100.

Weliky M, Bosking WH, Fitzpatrick D (1996) A systematic map of direction preference in primary visual cortex. Nature 379:725–728.

